# An End-to-End Deep Learning Speech Coding and Denoising Strategy for Cochlear Implants

**DOI:** 10.1101/2021.11.04.467324

**Authors:** Tom Gajecki, Waldo Nogueira

**Affiliations:** Department of Otolaryngology, Medical University Hannover and Cluster of Excellence Hearing4all, Hannover, 30625, Germany

**Keywords:** Cochlear Implant, Deep Learning, Sound Coding Strategy, Speech Enhancement

## Abstract

Cochlear implant (CI) users struggle to understand speech in noisy conditions. To address this problem, we propose a deep learning speech denoising sound coding strategy that estimates the CI electric stimulation patterns out of the raw audio data captured by the microphone, performing end-to-end CI processing. To estimate the relative denoising performance differences between various approaches, we compared this technique to a classic Wiener filter and to a conv-TasNet. Speech enhancement performance was assessed by means of signal-to-noise-ratio improvement and the short-time objective speech intelligibility measure. Additionally, 5 CI users were evaluated for speech intelligibility in noise to assess the potential benefits of each algorithm. Our results show that the proposed method is capable of replacing a CI sound coding strategy while preserving its general use for every listener and performing speech enhancement in noisy environments, without sacrificing algorithmic latency.

## 1. INTRODUCTION

A cochlear implant (CI) is a surgically implanted medical device that can restore hearing to a profoundly deaf person. In general, CI users achieve good speech intelligibility in quiet conditions. When compared to normal-hearing listeners, however, CI users need significantly higher signal-to-noise ratios (SNRs) to achieve the same speech intelligibility [1]. This fact motivates researchers to investigate different speech enhancement techniques to improve the SNR of the incoming signal in acoustically challenging conditions [2].

The CI sound coding strategy is responsible for computing the electric stimulation current levels from the audio captured by the CI sound processor microphone. It uses a filter bank that decomposes the incoming sound into different analysis sub-band signals, which are used to encode electric pulses to stimulate the auditory nerve.

Previous research has shown that single-channel noise reduction algorithms can be used as front-end processors prior to the sound coding strategy to improve speech intelligibly of CI users [3, 4]. Single-channel noise reduction algorithms generally convert the input signal into the spectral domain and apply masks to emphasize the frequency bands with high SNRs and attenuate the noisy ones, performing an enhancement of the target signal [5]. These algorithms rely on signal processing methods that include spectral subtraction such as Wiener filtering [6,7]. Currently, estimating accurate masks for CI users while minimizing distortions on speech signals still remains a challenge. In fact, Wiener filters, like the ones used in commercial CI sound processors, provide limited or no benefit under non-stationary noise conditions.

For non-stationary noisy backgrounds, speech enhancement can be achieved by means of spatial filtering algorithms (i.e., beamformers), assuming that the target speech and masking noise are spatially separated [5,8]. Nonetheless, more recently, data-driven approaches based on deep neural networks (DNNs), have been also successful at improving speech understanding in non-stationary background noise conditions for CI listeners [9,10]. These algorithms, however, perform front-end processing and are not well integrated into the CI sound coding strategy. In order to optimize speech enhancement for CIs, it may be beneficial to design algorithms that consider the CI processing scheme. Thus, there has been some work done specifically for CIs, where DNNs are included in the CI signal path [11,12]. These approaches perform noise reduction, for example, by directly applying masks in the filter bank used by the CI sound coding strategy. However, these approaches tend to rely on the spectrum of the sound or on other spectro-temporal features [11], limiting speech separation performance. Recently, several speech enhancement and audio source separation models that operate directly on time-domain audio signals have been proposed [13, 14, 15, 16, 17]. These end-to-end (audio-to-audio) approaches offer advantages, as fewer assumptions related to the magnitude and phase of the spectrum are required while obtaining high performance.

Here we propose a CI end-to-end (audio-to-electrodogram) speech coding and enhancement method that uses the audio captured by the microphone in the sound processor to estimate the levels at which the inserted electrodes should be mapped onto for electrical stimulation of the auditory nerve. This new approach is designed to completely bypass the CI sound coding strategy, providing the listener with signals as natural as the original sound coding strategy would, while performing speech denoising. This end-to-end CI strategy may outperform a front-end DNN in terms of speech enhancement, as the estimated electrodograms have a lower dynamic range, less amplitude resolution, lack phase information, and are more redundant than raw audio signals [18], and therefore, may be easier to model.

The organization of the manuscript is as follows: section 2 presents the methods and materials, section 3 the evaluation of the speech enhancement algorithms using objective instrumental measures, and speech intelligibility tests in CI users. Section 4 presents the results and we conclude the manuscript in Section 4.

## 2. METHODS & MATERIALS

### 2.1. Algorithms

#### Advanced combination encoder (ACE)

The acoustic signal is first captured by the CI microphone and sampled at 16 kHz. Then, a filter bank implemented as a 128 point fast Fourier transform (FFT), commonly with a 32 point hop size, is applied, introducing a 2 ms algorithmic latency (this will depend on the stimulation rate; CSR). Next, an estimation of the desired envelope is calculated for each spectral band *E*_*k*_, (*k* = 1, …, *M*), each of which is allocated to a single electrode, representing one channel. Out of the *M* channels contained in each audio frame, only the *N* most energetic ones are selected for stimulation. Typical values for *M* and *N* are 22 and 8, respectively. The selected bands are subsequently non-linearly compressed by a loudness growth function (LGF) given by:

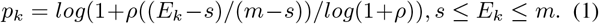

For values of *E*_*k*_ below base level *s, p*_*k*_ is set to 0, and for values of *E*_*k*_ above saturation level *m, p*_*k*_ is set to 1. For a detailed description of the parameters *s, m* and *ρ*, refer to [19]. Finally, the last stage of the sound coding strategy maps *p*_*k*_ into the subject’s dynamic range between threshold levels (THLs) and most comfortable levels (MCLs) for electrical stimulation. For each audio frame, the *N* selected electrodes are stimulated sequentially, representing one stimulation cycle. The number of cycles per second thus determines the CSR. A block diagram representing the previously described processes is shown in Figure 1a; ACE. The graphical representation of the current applied to each electrode over time is known as an electrodogram (Figure 2).

**Fig. 1.**
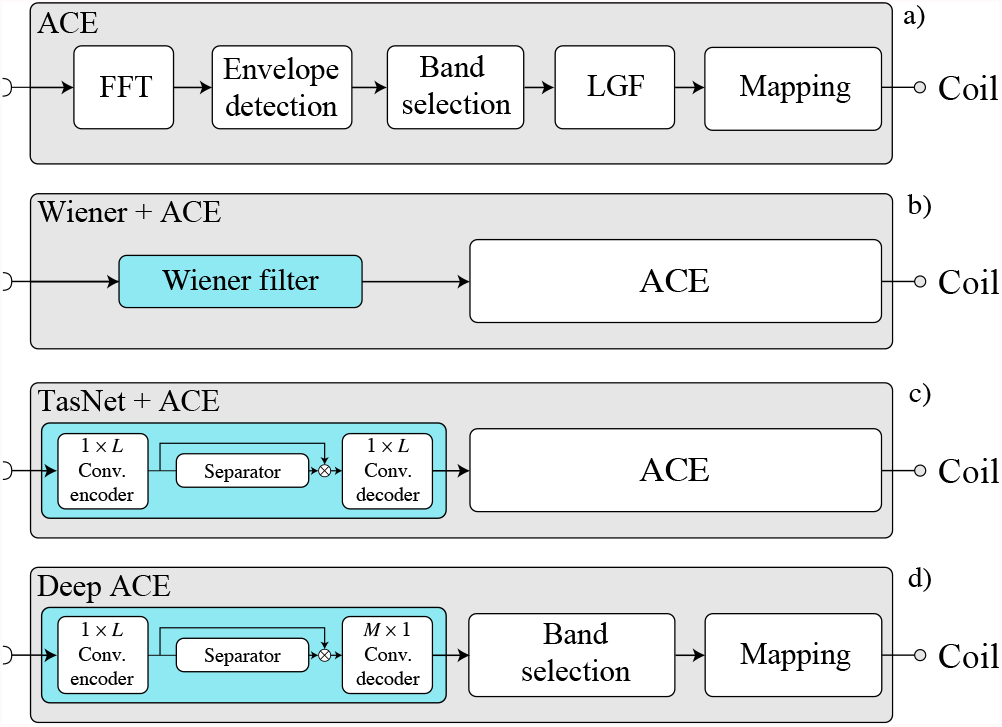
Block diagrams of the four different signal processing systems. In c) and d) *L* refers to the *length of the filters* used in the encoder and decoder (refer to Table 1).

**Fig. 2.**
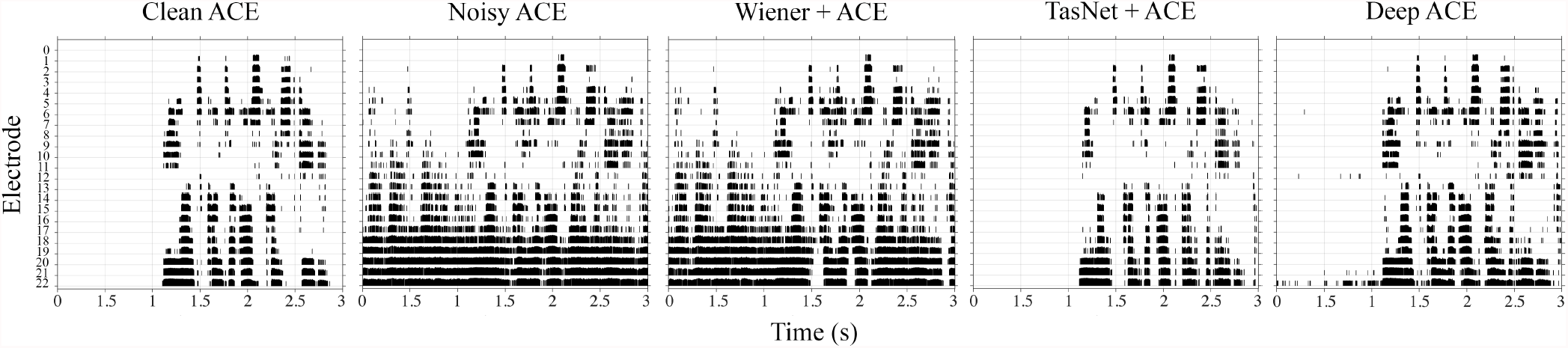
Electrodograms for the clean and noisy speech produced by ACE (first two electrodograms) and the electrodograms produced by the enhancing algorithms.

#### Baseline speech denoising algorithm (Wiener)

Here, we use a classic front-end signal processing method based on Wiener filtering, a widely used technique for speech denoising that relies on a priori SNR estimation [7] (Figure 1b; Wiener+ACE). This algorithm is used in all commercially available single channel noise reduction systems included in CIs [20, 21]. Therefore, this classic algorithm is an appropriate baseline to use when developing new speech enhancement methods in the context of CIs [11].

**Table 1.** Hyper-parameters used for training the models.

| Description | Value |
| --- | --- |
| Number of filters in the autoencoder | 64 |
| Length of the filters | 32 |
| Number of channels in the bottleneck blocks | 64 |
| Number of channels in the skip-connections | 32 |
| Number of channels in the convolutional blocks | 128 |
| Kernel size in the convolutional blocks | 128 |
| Number of convolutional blocks in each repeat | 3 |
| Number of repeats | 2 |

#### Baseline deep learning speech denoising algorithm (TasNet)

The DNN based baseline system used in this study is the well-known conv-TasNet (which will we refer to as “TasNet” for simplicity) [13]. This algorithm performs end-to-end audio speech enhancement and feeds the denoised signal to ACE (Figure 1c; TasNet+ACE). The TasNet structure has proven to be highly successful for single-speaker speech enhancement tasks, improving state-of-the-art algorithms, which is the main reason that it is commonly used as a baseline model [17]. The hyper-parameters chosen for the TasNet baseline are shown in Table 1. Note that the filter length at the encoder causes an algorithmic latency of 2 ms, which together with ACE results in a total algorithmic latency of 4 ms.

#### End-to-end sound coding strategy for CIs (Deep ACE)

Here we propose a new strategy that combines the ACE with the structure of TasNet [13]. Deep ACE takes the raw audio input captured by the microphone and estimates the output of the LGF (Figure 1d; Deep ACE). By predicting *p*_*k*_ ∈ [0, 1], the strategy is not only independent of individual CI fitting parameters, but it also retains the 2 ms total algorithmic delay introduced by the standard ACE strategy. The enhancer module in deep ACE is similar to the one in TasNet+ACE, differing only in the activation function used in the encoder and in the output dimensionality of the decoder (Figure 1c and d).

The activation function used in deep ACE encoder is given by:

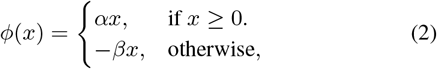

where {(*α, β*) ∈ ℝ^0+^ × ℝ^0+^} are trainable scalars that control the positive and negative slope of the rectifier. This activation function guarantees that the coded signal is represented by real positive values (for nonzero input values). The other difference between the enhancer blocks of TasNet+ACE and deep ACE is related to the output dimensionality. The TasNet enhancer module estimates an output in the time domain, every temporal convolutional window, whereas deep ACE will estimate the LGF output in the CI channel domain, ready to perform band selection. The code for training and evaluating deep ACE can be found online^1^.

### 2.2. Datasets

#### Dataset 1

The audio dataset was provided by the 1st Clarity enhancement challenge [22]. It consists of 6,000 scenes including 24 different speakers. The development dataset, used to monitor the model performance during training, consists of 2,500 scenes including 10 target speakers. Each scene corresponds to a unique target utterance and a unique segment of noise from an interferer, mixed at SNRs ranging from -6 to 6 dB. The two sets are balanced for the target speaker’s gender. Binaural room impulse responses (BRIRs) were used to model a listener in a realistic acoustic environment. The audio signals for the scenes are generated by convolving source signals with the BRIRs and summing. BRIRs were generated for hearing aids located in each listening side, providing 3 channels each (front, mid, rear). From which only the front microphone was used.

#### Dataset 2

In addition to dataset 1, the Hochmair, Schulz, Moser (HSM) sentence test [23], composed of 30 lists with 20 everyday sentences each (106 words per list) was used. The HSM sentences were mixed with interfering multiple-speaker-modulated speech-weighted noise source (ICRA7) [24] and interfering Consultatif International Téléphonique et Télégraphique (CCITT) noise [25], at SNRs ranging from -5 to 5 dB. Speech and noise signals were convolved with a BRIR [26] and presented in a virtual acoustic scenario at a distance of 80 cm in front of the listener.

#### Train, validation and test datasets

All data were downsampled to 16 kHz. To train the models, the training set of dataset 1 was mixed with 30% of dataset 2. To validate and optimize the models, the validation set of dataset 1 was used. Lastly, for final testing, the remaining 70% of dataset 2 was used.

### 2.3. Model Training

The models were trained for a maximum of 100 epochs on batches of two 4-second long audio segments captured by a single CI. The initial learning rate was set to 1e-3. The learning rate was halved if the accuracy of the validation set did not improve during 3 consecutive epochs, early stopping with a patience of 5 epochs was applied as a regularization method, and only the best performing model was saved. For optimization, Adam [27] was used to optimize the desired cost function, which depended on the algorithm to be trained.

#### TasNet+ACE cost function

In the case of the TasNet+ACE algorithm, the optimizer was used to maximize the scale-invariant (SI) SNR [28] at the output of the TasNet. The SI-SNR between a given signal with *T* samples, ***x*** *∈* ℝ^1×*T*^ and its estimate 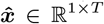 is defined as:

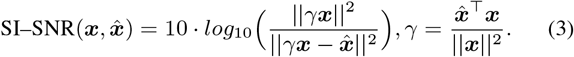

#### Deep ACE cost function

Because the enhancer module will estimate the output at the LGF of ACE, the optimizer will be used to minimize the mean-squared-error (MSE) between the predicted and target signals. The MSE across electrodes between an *F* –frame target signal, ***p*** *∈* ℝ^*M×F*^ and its estimate 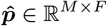, is defined as:

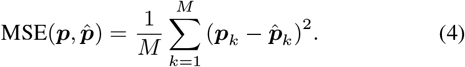

#### Hyper-parameter optimization

To assess which model size was the best to train the algorithms, we factorized the problem by examining the effect on the validation error as a function of the skip connection size. We performed 5 independent training sessions for different skip connection channel sizes {4, 8, 16, 32, 128, 256, 512, 1024}. The model with the lowest validation error was chosen for the final evaluation.

Table 1 shows the used hyper-parameters of the implemented models. For a detailed description of these hyper-parameters refer to [13].

The models were trained and evaluated using a PC with an Intel(R) Xeon(R) W-2145 CPU @ 3.70GHz, 256 GB of RAM, and an NVIDIA TITAN RTX as the accelerated processing unit.

## 3. EVALUATION

### 3.1. Objective Instrumental Evaluation

#### SNR Improvement

For a given algorithm, the SNR (eq. 3 with *γ* = 1) improvement with respect to the unprocessed signal (noisy ACE) will be reported. This is simply computed as follows: SNRi = SNR_proc._ *−* SNR_unproc._. To obtain the processed signals in the time domain for each of the algorithms, the generated electrodograms were resynthesized using a sine vocoder with a THL of 100 and an MCL of 150 clinical units (refer to [19]). Then, equation 3 was applied to the corresponding vocoded signals.

#### STOI

This measure is used to predict the speech intelligibility performance for each of the tested algorithms. To compute it, the generated electrodograms were resynthesized using the vocoder described in the previous subsection. The original noiseless, clean speech signals served as reference signals (raw speech signals captured by the microphone). The resynthesized audio waveforms and the reference signals were used to obtain the short-time objective intelligibility (STOI) measure [29].

### 3.2. Listening Evaluation

#### Participants

5 postlingually deafened CI users participated in the study. All participants were native German speakers and traveled to the Hannover Medical School (MHH) for a 2-hour listening test and their travel costs were covered. The experiment was granted with ethical approval by the MHH ethics commission. A synopsis of the pertinent patient-related data is shown in Table 2.

**Table 2.**
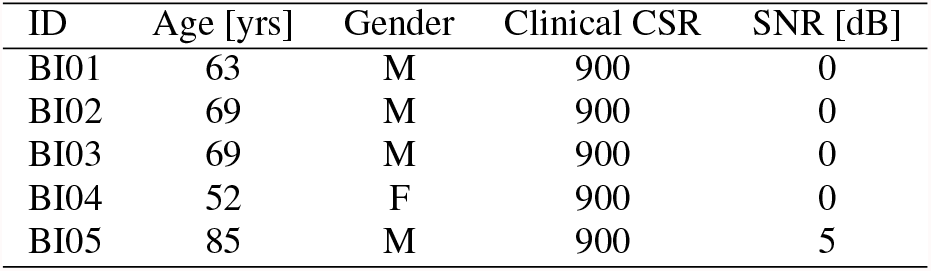
Listener demographics and etiology. The clinical CSR expressed in pulses per second (pps) is the one that participants were using in their clinical speech processors. The last column indicates the SNR at which each subject was tested.

#### Test scenario

For the listening experiments in CIs, the remaining 70% of the test dataset, mixed with ICRA7 noise, was used. Stimuli were delivered via direct stimulation through the RF GeneratorXS interface (Cochlear Ltd., Sydney, Australia) with MATLAB (Mathworks, Natick, MA) via the Nucleus Implant Communicator V.3 (Cochlear Ltd., Sydney, Australia). The CSR used in this study to train and evaluate the models was 1000 pps. Speech intelligibility in noise was measured by means of the HSM sentence test [23]. Subjects were asked to repeat sentences out loud as accurately as possible. Each listening condition was tested twice with different sentence lists, then the final score was computed by taking the mean number of correct words for each condition. The conditions were blinded to the subjects.

## 4. RESULTS

### 4.1. Objective Evaluation Results

#### SNR improvement

Figure 3 shows the SNRi obtained by the investigated algorithms w.r.t. ACE, in two different background noises and three input SNRs.

**Fig. 3.**
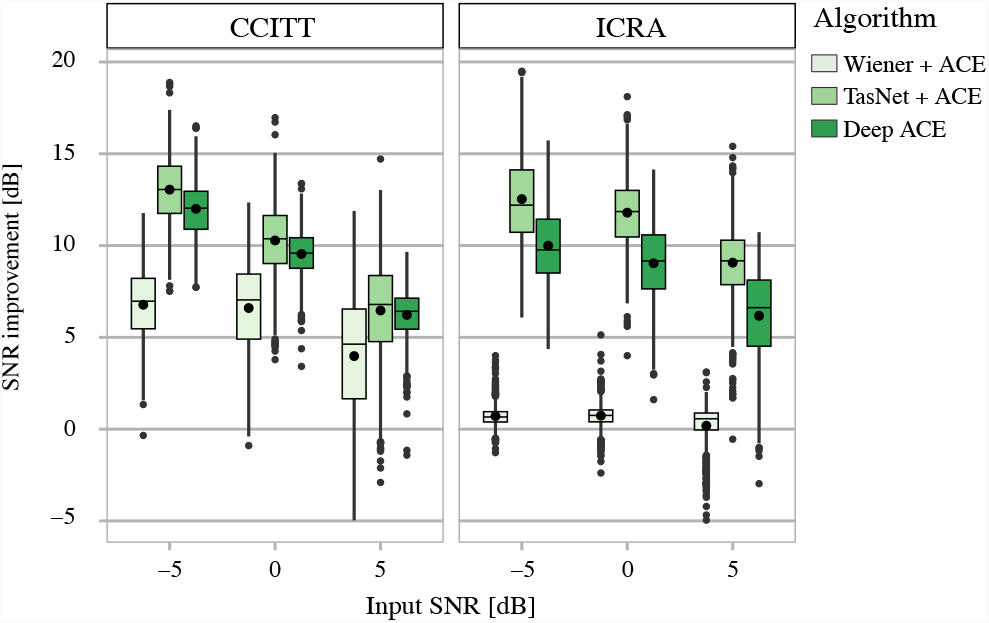
SNR improvement in dB for the tested algorithms in CCITT noise and in ICRA noise for the different SNRs.

#### STOI

The mean STOI scores obtained by the ACE sound coding strategy, TasNet+ACE, and deep ACE for speech signals *without interfering noise* were 0.8, 0.79, and 0.78, respectively.

Figure 4 shows the STOI results obtained by the tested algorithms, for two different background noise types and three input SNRs.

**Fig. 4.**
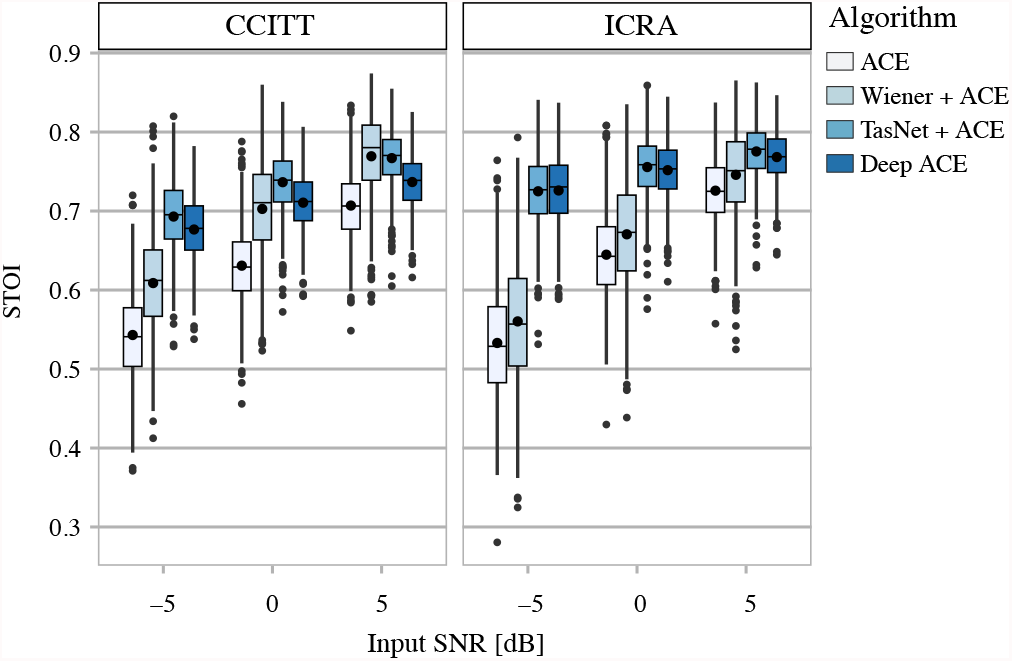
STOI scores obtained by the tested algorithms in CCITT noise and in ICRA noise for the different SNRs.

### 4.2. Listening Results

Figure 5 shows the percentage of understood words in quiet and mixed with ICRA7 noise at an input SNR indicated in Table 2.

**Fig. 5.**
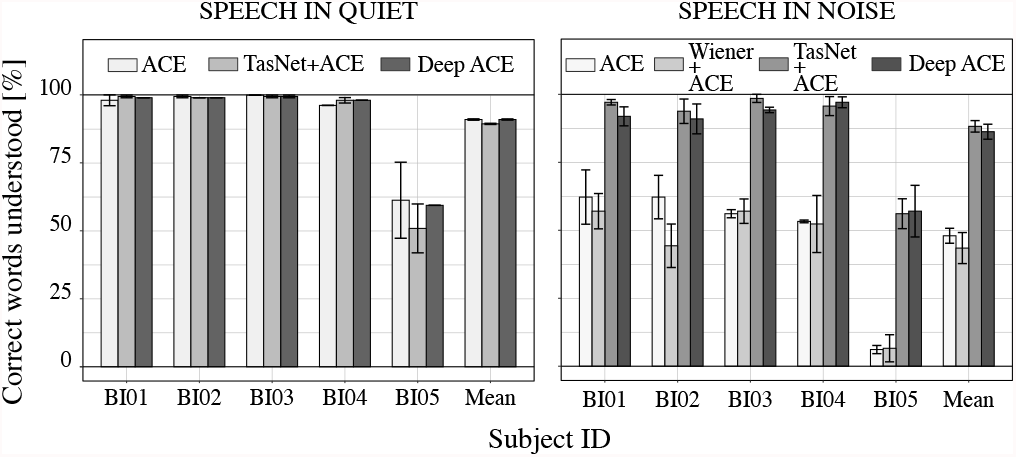
Individual and mean percentage of correct understood words by subject for the HSM sentence test in quiet and in ICRA7 noise.

## 5. CONCLUSIONS

In this work, we have presented an adaptation of the TasNet model for speech denoising to a CI sound coding strategy; deep ACE. This approach allows reducing the processing complexity of the ACE sound coding strategy while performing noise reduction for CIs. We found that the proposed method and a front-end speech enhancement method based on TasNet do not affect speech understanding in quiet when compared to ACE. In the context of speech enhancement, deep ACE showed slightly worse objective performance than the front-end TasNet approach. This may be potentially related to a sub-optimal cost function used to minimize the error between the input and target electric stimulation patterns. However, the speech perception scores obtained with deep ACE and the front-end TasNet, were very similar. It is important to remember that deep ACE reduces the algorithmic latency with respect to the front-end TasNet by 2 ms (introduced by the ACE sound coding strategy). The proposed method has the potential to completely replace any CI sound coding strategy while keeping its general usage for every listener and performing speech enhancement in noisy conditions.

## Footnotes

1 https://github.com/APGDHZ/DeepACE

